# EnrichMet: R Package for Integrated Pathway and Network Analysis for Metabolomics

**DOI:** 10.1101/2025.08.28.672951

**Authors:** Yonatan Ayalew Mekonnen, Neha Dhake, Vanessa Rubio, Shreya Jaiswal, Isis Narváez-Bandera, Ashley Lui, Augustine Takyi, Hayley Ackerman, John Koomen, Elsa Flores, Paul A. Stewart

## Abstract

Advances in metabolomics have significantly improved our understanding of cellular processes by enabling the identification of hundreds of metabolites in a single experiment. These developments provide valuable insights into complex metabolic networks. While efforts have been made to develop pathway enrichment analysis (PEA), existing implementation often require multiple steps, rely on web-based interfaces, or depend on R packages configuration that may affect reproducibility and ease of use. To overcome these limitations, we introduce EnrichMet, an R package for fast, flexible, and reproducible pathway enrichment analysis. EnrichMet modules support over-representation analysis of pathways, metabolite set enrichment analysis (MetSEA), and network-based pathway analysis. The package streamlines the workflow by combining curated pathway information from the Kyoto Encyclopedia of Genes and Genomics (KEGG) and employs Fisher’s Exact Test to identify significantly enriched pathways. Benchmark analyses show that enrichment on sample data completes in approximately 3 seconds. EnrichMet offers both a command-line and a user-friendly Shiny interface, enabling accessibility for users with or without programming experience. Through case studies on experimental metabolomics datasets, we demonstrated that EnrichMet delivers accurate and comprehensive pathway enrichment results while minimizing computational time and simplifying user interaction. Furthermore, its flexible framework supports extensions to other data types and knowledge bases beyond KEGG, as illustrated through a lipidomics case study. By unifying performance, reproducibility, usability, and visualization within a single package, EnrichMet facilitates deeper insights and promotes efficient, transparent, and reproducible research practices.

**Availability and implementation:** (https://github.com/biodatalab/enrichmet.git)

## Introduction

High-throughput metabolite identification in a single experiment reveals unprecedented insights into the complexity of cellular processes. However, interpreting these metabolites in a meaningful biological context remains challenging [1]. In metabolomics, a pathway typically represents a set of analytes, such as metabolites, enzymes, or reactions, that participate in related biological processes. Pathway Enrichment Analysis (PEA) [2-4] and Metabolite Set Enrichment Analysis (MetSEA) [5] address this challenge by statistically evaluating whether metabolites associated with particular pathways occur more frequently than expected by chance. These approaches identify groups of related features that may be altered under specific experimental conditions, helping researchers move from lists of detected metabolites to interpretable biological hypotheses [4]. Building on this concept, our tool provides an intuitive and streamlined framework for enrichment analysis, helping users uncover functional patterns and derive biologically meaningful insights from metabolomics data with greater confidence.

Several tools have been developed for pathway enrichment in metabolomics, including MetaboAnalyst [6], Mummichog [7], PIUMet [8], PUMA [9], and FELLA [10], with MetaboAnalyst being among the most widely used platforms [11]. These tools have advanced the field by enabling pathway-level interpretation of metabolomics data, but many require multiple installation steps, external dependencies, or web interfaces that can limit reproducibility and flexibility. A comparative summary of the software tools is provided in Table 2. To address these limitations, we developed EnrichMet, an R package that performs pathway enrichment in seconds, can be installed with a single command, and runs entirely locally. The tool is designed for both accessibility and extensibility, allowing users to perform analyses quickly while retaining the flexibility to adapt the framework for other data types, species, or pathway databases.

### Performance and Implementation

EnrichMet demonstrated a marked improvement in computational efficiency compared to existing tools such as MetaboAnalystR and FELLA. Using an Apple MacBook Pro equipped with an M3 Pro chip and 36 GM of memory running macOS version 15.7.1, EnrichMet completed the pathway analysis for representative dataset of 94 metabolites in approximately 3 seconds, compared with approximately 28 seconds for MetaboAnalystR and approximately 14 minutes for FELLA. This substantial performance advantage makes EnrichMet particularly ideal for high-throughput or iterative analyses, where speed is critical. EnrichMet offers full transparency and control over the analysis process, allowing users to easily access results and integrate the package into larger R workflows. Beyond performance, EnrichMet offers full transparency and control over the analytical process. All computations are executed entirely within the R environment, ensuring reproducibility, customization, and seamless integration into broader data analysis pipelines.

### Modular Framework

EnrichMet is implemented as a fully modular framework that can be executed through a single functional call. Each internal function implemented as a modular component accompanied illustrative examples demonstrating its functionality. When the main EnrichMet function is invoked, it sequentially executes all underlying modules, providing seamless workflow. This modular design ensures transparency, ease of use, and flexibility, enabling users to easily modify or manage the output.

### EnrichMet: integrated pathway and network analysis

EnrichMet leverages curated pathway data from the Kyoto Encyclopedia of Genes and Genomes (KEGG) and features modular analysis components that enable over-representation analysis, metabolite set enrichment analysis (MetSEA), and topology-based network analysis. By integrating metabolite input and background datasets, EnrichMet identifies enriched pathways with minimal user intervention, often requiring only a single function call. The full workflow is illustrated in Figure 1. This manuscript highlights EnrichMet’s streamlined design, statistical rigor, and adaptability, demonstrating its utility for metabolomics data analysis through a case study dataset.

**Figure 1.**
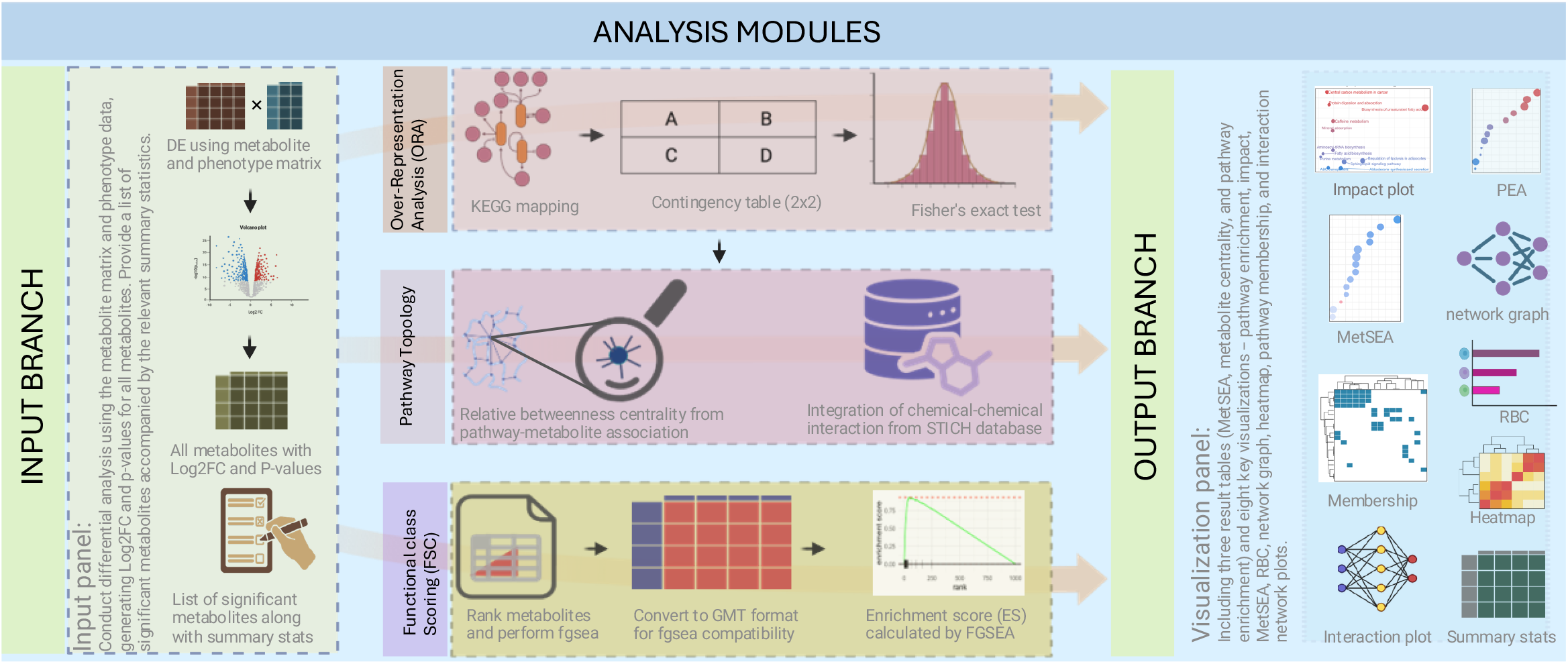
Overview of the EnrichMet data analysis workflow.

### Overrepresentation analysis functionality

Metabolite and pathway association data were obtained from the Kyoto Encyclopedia of Genes and Genomes (KEGG) database. The analysis utilized two dataset sources: an input list of differentially expressed metabolites identified based on statistical significance (P-value) and magnitude of change (fold-change), and a predefined background set consisting of all known KEGG metabolites annotated to KEGG pathways, obtained via the KEGGREST R package. For convenience, background sets for human, mouse, yeast, fruit fly and zebrafish are provided although users may generate or supply background data for other species of interest. These datasets were organized to map each pathway to its corresponding metabolites, providing a robust framework for statistical analysis. For each pathway, a 2x2 contingency table was constructed as shown in Table 2. Fisher’s Exact Test [12] was applied to the contingency table to assess whether the number of identified metabolites in a pathway significantly exceeded expectations. The output of Fisher’s Exact Test is a p-value, which indicates the likelihood that the observed enrichment or depletion of metabolites in a pathway is due to random chance. To account for multiple comparisons and enhance the reliability of statistical significance, p-values were adjusted using the Benjamini-Hochberg method and Q-values were also computed to provide an additional measure of significance. The final output ranks pathways by significance, highlighting those enriched with the identified metabolites.

The p value from Fisher’s Exact Test is calculated as [12]:

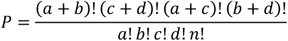

Where, a, b, c, d are counts from the contingency table (Table 1).

**Table 1.** Comparative overview of software tools used in metabolomics pathway enrichment analysis.

| Tools | Methods | Input data | Pathway Topology | Visualization Options | R package | Shiny App | Web Interface | Fast for Large Data | Independent of DB Building | Yes Score |
| --- | --- | --- | --- | --- | --- | --- | --- | --- | --- | --- |
| EnrichMet | Fisher’s exact test, fgsea | Identified metabolites | yes | yes | yes | Yes | no | yes | yes | 86% |
| MetaboAnalystR | Fishers’ exact test, hypergeometric test, GSEA, | Identified metabolites | yes | Moderate | yes | no | yes | moderate | yes | 57% |
| Mummichog | Permutation-based enrichment | m/z values | yes | limited | no | no | no | yes | yes | 43% |
| FELLA | Diffusion over KEGG graph, hypergeometric test | Identified metabolites | yes | limited | yes | no | no | no | no | 29% |
| PIUMet | Bayesian graphical model | m/z, RT | yes | no | no | no | yes | no | no | 29% |
| PUMA | Network-based scoring | Metabolite, gene, protein data | yes | no | no | no | no | no | no | 14% |

**Table 2.**
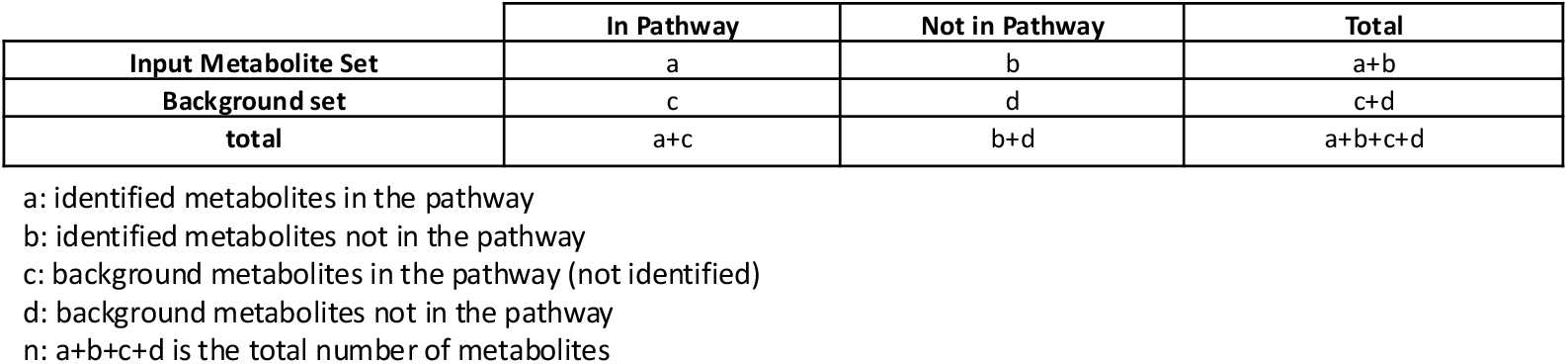
Contingency table used for Fisher exact test in pathway enrichment analysis.

*n*= a+b+c+d is the total number of metabolites.

Results from the enrichment analysis were summarized in tabular format and visualized using dot plots of -log10(adjusted p-values), highlighting the top significantly enriched pathways. To further assess its performance, results were compared against those generated using MetaboAnalystR and FELLA R packages, demonstrating consistency in findings while showcasing a more streamlined and reproducible analytical process.

### Metabolite set enrichment analysis (MetSEA) functionality

Pathway enrichment analysis typically relies on predefined significance thresholds to select metabolites or feature, which can be arbitrary and introduce bias [13]. In contrast MetSEA leverages the entire ranked list of metabolites based on the fold change and p-values, without requiring cutoffs. This approach enables the detection of coordinated and subtle changes across pathways, offering a more systematic and less biased assessment of biological enrichment [5]. EnrichMet also performs metabolite set enrichment analysis (MetSEA) by integrating the fgsea R package, which implements a computationally efficient algorithm for Gene Set Enrichment Analysis (GSEA). Pathway annotations are converted into a Gene Matrix Transposed (GMT) format to ensure compatibility with fgsea. Metabolites are ranked based on their differential expression metrics (log2 fold change and p-values), creating a ranked list required for fgsea [14].

The enrichment Score (ES) represents the maximum deviation from zero observed as the algorithm walks down the ranked list of the metabolites. The normalized enrichment score (NES) is subsequently derived by scaling the ES against the distribution of ES values obtained from permutation testing, thereby accounting for differences in the pathway size and variability [14]. The analysis outputs significant pathways, their adjusted p-values, and enrichment statistics. A positive or negative NES accompanied by small adjusted p-value (< 0.05) indicates that the corresponding pathway is significantly enriched. A positive NES reflects a coordinated increase (upregulation) in the abundances of metabolites within that pathway, whereas a negative NES indicates a coordinated decrease (downregulation) relative to the reference condition. These results are stored as an S3 data frame, a standardized R structure that enables convenient access and downstream analyses, offering a comprehensive summary of enriched pathways and their associated metrics.

### Topological measure functionality

Network analysis provides a framework for characterizing the complex interactions among metabolites within biological systems. In this approach, metabolites are represented as nodes and their shared pathway as edges, thereby capturing topological relationships. Within this framework, betweenness centrality (BC) measures the frequency of which a metabolite lies on the shortest paths connecting other metabolites, reflecting its role in mediating network communication. Therefore, metabolites with high BC serve as a central connector that may influence multiple pathways, whereas those with low BC are more isolated. Centrality metrics were calculated using the igraph package in R, and hub metabolites were identified based on their connectivity, defined as the number of edges linking each metabolite to others within the network. Betweenness centrality is calculated as [15]:

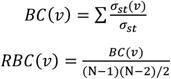

where σ_*st*_ is the total number of shortest paths between nodes *s* and *t*, σ_*st*_(*v*) is the number of those paths that pass-through node *v*. To normalize BC, we compute relative betweenness centrality (*RBC*) which is normalized form of BC and N is the total number of nodes in the network.

To account the relative importance of pathways in a global metabolic network, we calculated the pathway impact score which quantifies the extent of which a pathway contributes to overall network connectivity and information flow. The pathway impact score reflects not only the number of metabolites involved but also their structural influence within the network. The pathway impact score was calculated as:

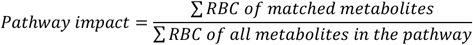

In addition to pathway-based networks derived from KEGG, we construct a metabolite-metabolite interaction network using the known chemical-chemical relationships from STITCH database to capture biological meaningful associations between metabolites beyond predefined pathway boundaries. While KEGG defines functional association through shared pathway membership, STITCH integrated evidence from multiple sources including experimental data, literature co-occurrence, and structural similarity to infer potential biochemical or functional interaction [16]. This approach provides a complementary context by highlighting metabolites that may co-function or participate in related biochemical processes even if they are not annotated within the same KEGG pathway. To ensure biological relevance, STITCH interactions were filtered based on combined score (≥100).

### Case study of control vs mutant lung cancer

This study used a dataset generated from metabolomic profiles comparing control and mutant lung cancer. The EnrichMet workflow can perform differential analysis through the running run_de function by providing the appropriate input object (Figure. S1). If summary statistics are already generated, then can be supplied directly to the EnrichMet workflows for downstream analysis. We selected significant metabolites (absolute fold change ≥ 1 and adjusted p value < 0.05) to apply Fisher’s Exact Test for pathway enrichment analysis. Significant pathways were obtained after adjusting for multiple comparisons using the Benjamini-Hochberg method (adjusted p-value < 0.05). The dot plot illustrates the -log10(adjusted p-values) of the most enriched pathways, highlighting key biological processes in Figure 2A. The pathway impact plot shows prioritizing pathways that are not only statistically enriched but also structurally critical in the biological networks. For example, Figure 2B indicates that pyrimidine metabolism appears as the top pathway based on its adjusted p value. However, the caffeine metabolism is also significant and shows a strong pathway impact, which may justify prioritizing this pathway over pyrimidine metabolism. Therefore, the pathway enrichment and impact analysis in alignment with the MetSEA results (Figure 2C), identifies two main categories: metabolic reprogramming and energy production, nutrient uptake and homeostasis. The metabolites involved in these pathways are visualized in Figures 3A and 3B.

**Figure 2.**
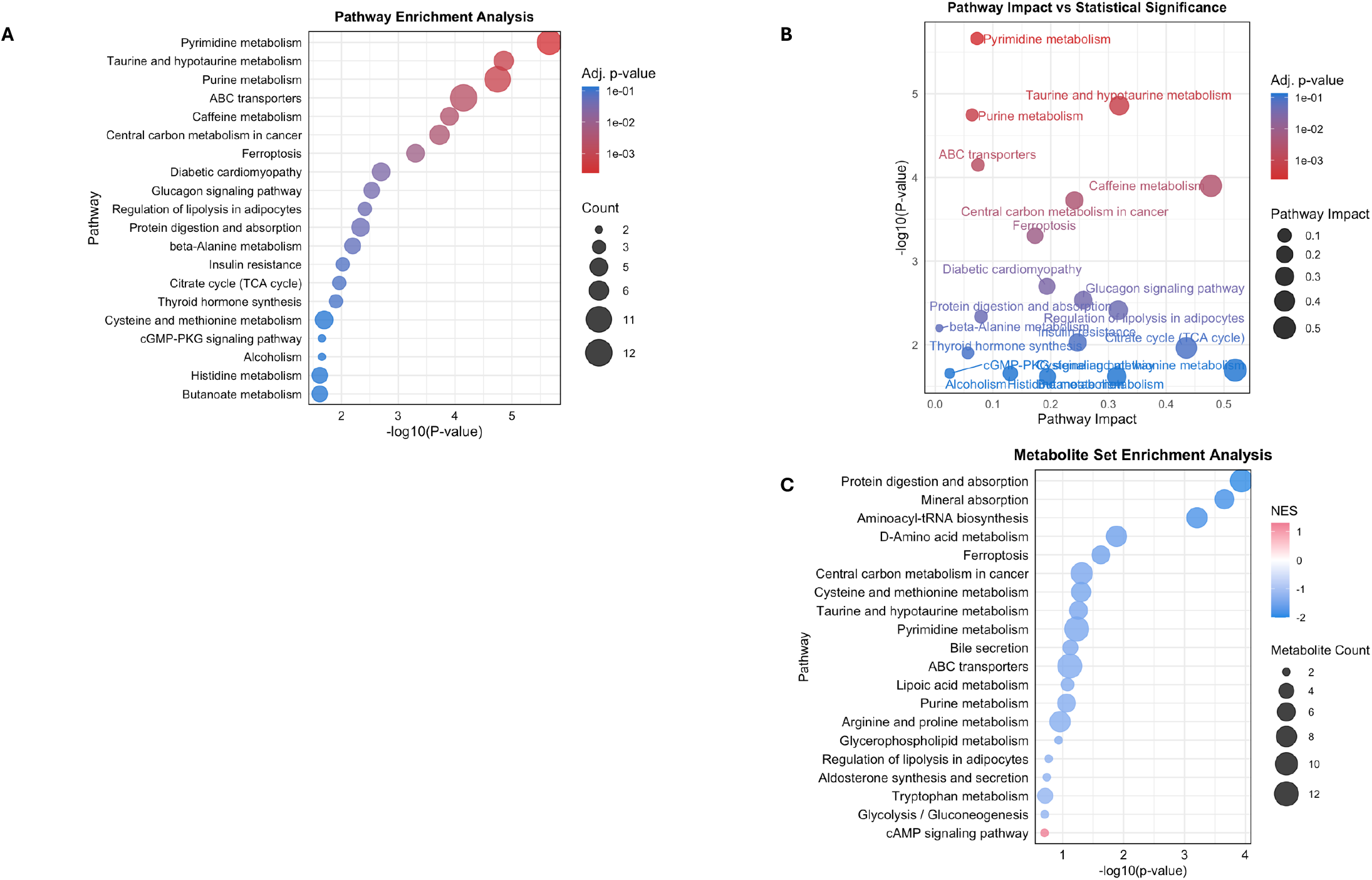
Overview of pathway enrichment and topology-based analysis results. **A)**. Pathway enrichment analysis displaying pathways ranked by –log_10_(p-value), calculated using Fisher’s exact test based on input metabolite-to-pathway mappings. **B)**. Pathway impact plot integrating enrichment significance with topological information, where Relative Betweenness Centrality (RBC) highlights the most topologically influential pathways. **C)**. Metabolite Set Enrichment Analysis (MetSEA), showing significantly enriched metabolite sets derived from ranked metabolite data.

**Figure 3.**
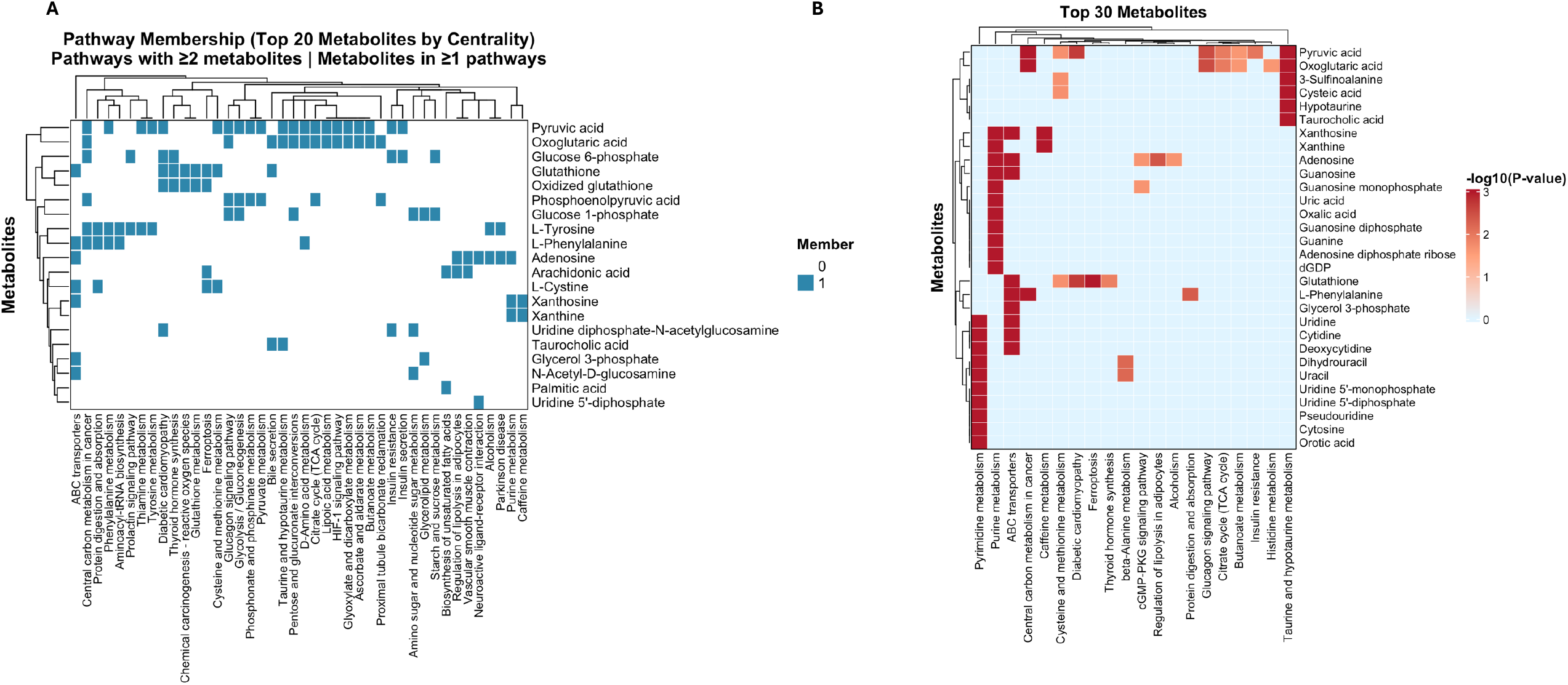
Visualization of pathway enrichment results and metabolite associations. **A)**. Pathway-metabolite association plot illustrating each pathway alongside its corresponding mapped metabolites. **B)**. Heatmap showing a gradient of –log_10_(p-values), representing the significance of enriched pathways. Only pathways with at least two mapped metabolites are included.

Notably, the network analysis identified hub metabolites including pyruvic acid (Figure 4A, B), Arachidonic acid (Figure 4A, B, C), and L-Tyrosine (Figure 4A, B). Moreover, the chemical-chemical interaction plot (Figure 4C) identifies Xanthine as a hub metabolite providing biological insights. Furthermore, recent studies have highlighted the potential roles of Cortexolone, Azelaic acid, and Uric acid in cancer biology [27-29].

**Figure 4.**
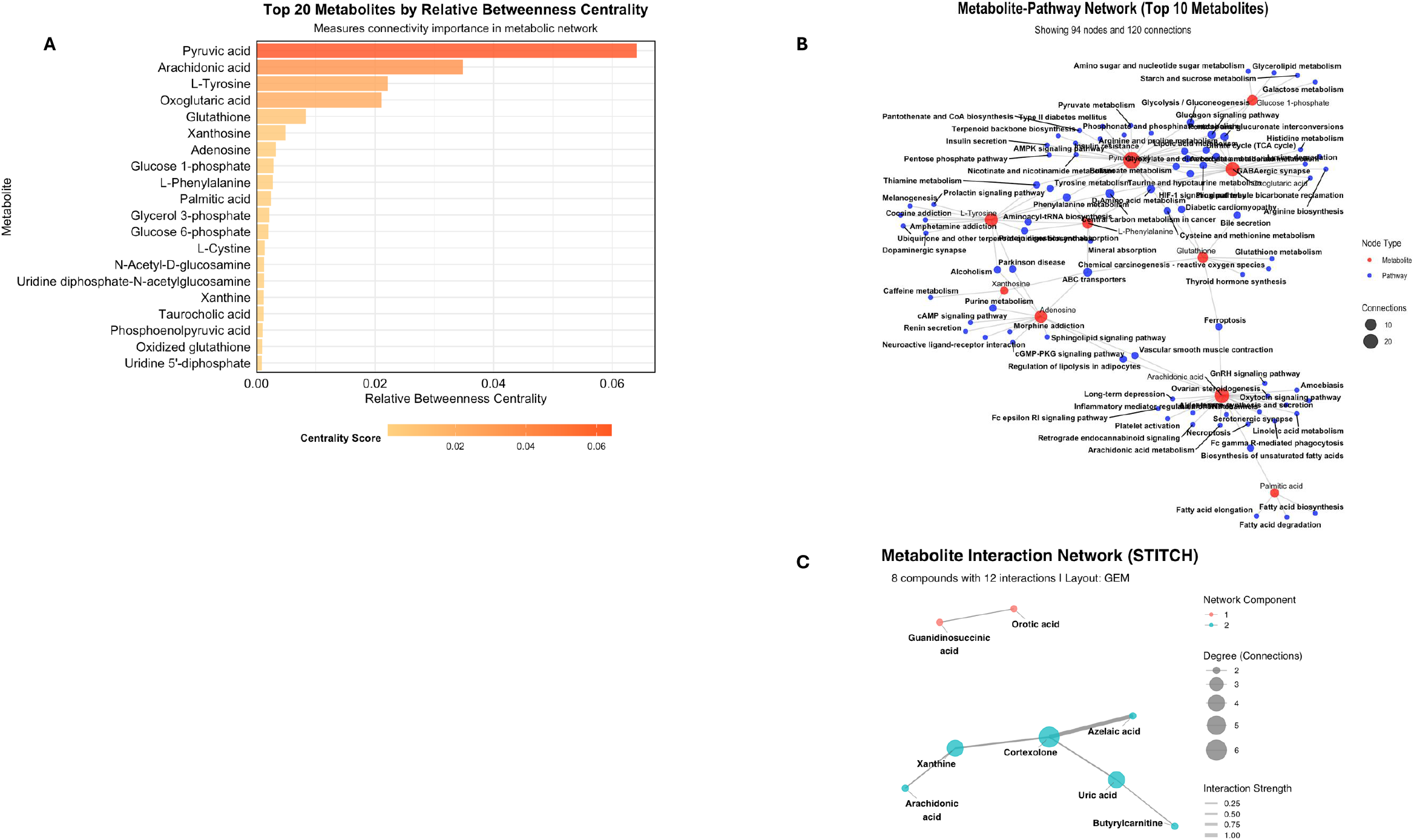
Network-based visualization of metabolite importance and interactions. **A)**. Relative betweenness centrality of input metabolites, reflecting their topological importance and potential regulatory roles within the metabolic network. **B)**. Metabolite-pathway interaction network, illustrating the connectivity between pathways and input metabolites and highlighting their functional associations. The size of the hub metabolites is corresponded to the value of RBC, edges are pathways and nodes are metabolites. **C)**. Chemical-chemical interaction network derived from STITCH. Metabolites are shown as nodes whose size scales with degree (the number of direct interaction partners). Edges represent experimentally validated or predicted biochemical relationship; and edge opacity is mapped to the rescaled STITCH combined score (weight), with darker edges indicating higher-confidence interaction.

### Showcasing Analytical Flexibility: Application to Lipidomics

To demonstrate the enrichment capabilities of EnrichMet, we applied our tool to a lipidomics dataset comparing samples harboring the control and mutant lung cancer. Lipid–ontology associations of this analysis were sourced from the LION database [30]. As illustrated in Figure S2, this example highlights the flexibility and robustness of our approach and underscores its potential applicability beyond metabolomics.

### Benchmarking

#### Streamlined Workflow

MetaboAnalyst is widely used in metabolomics community for many aspects of data analysis. However, installation of the package can be hindered by outdated or missing dependencies, particularly for packages not available in standard R repositories, and it remains uncertain whether MetaboAnalystR still depends on deprecated components requiring manual compilation. Similarly, the FELLA package requires building a local KEGG graph database, which is memory intensive and time consuming. This can significantly slow down performance, making FELLA less practical for high-throughput metabolomics datasets. However, EnrichMet offers a streamlined alternative for pathway enrichment analysis by performing the entire process with a single R function call. This one-step workflow reduces manual steps such as data formatting, and parameter selection minimizing user intervention, reducing potential errors and improving reproducibility. A shiny application of EnrichMET has also been developed to provide users with an intuitive and user-friendly interface, enabling efficient visualization, exploration, and interpretation of analysis results.

### Summary

EnrichMet streamlines pathway enrichment analysis, improving both computational efficiency and interpretability. The workflow allows enrichment and topological analysis to be performed in a single function call while also supporting modular, individual analyses. The versatile and user-friendly design enables researchers to prioritize biological insights over technical complexity. Combined with robust statistical methodologies, EnrichMet serve as a powerful tool for metabolomics and other omics-based pathway analyses. By integrating PEA, topological measures, and MetSEA, EnrichMet offers a comprehensive and robust statistical rigor, providing reliable results for comparison.

