## Supplemental Figures for "EnrichMet: R Package for Integrated Pathway and Network Analysis for Metabolomics"

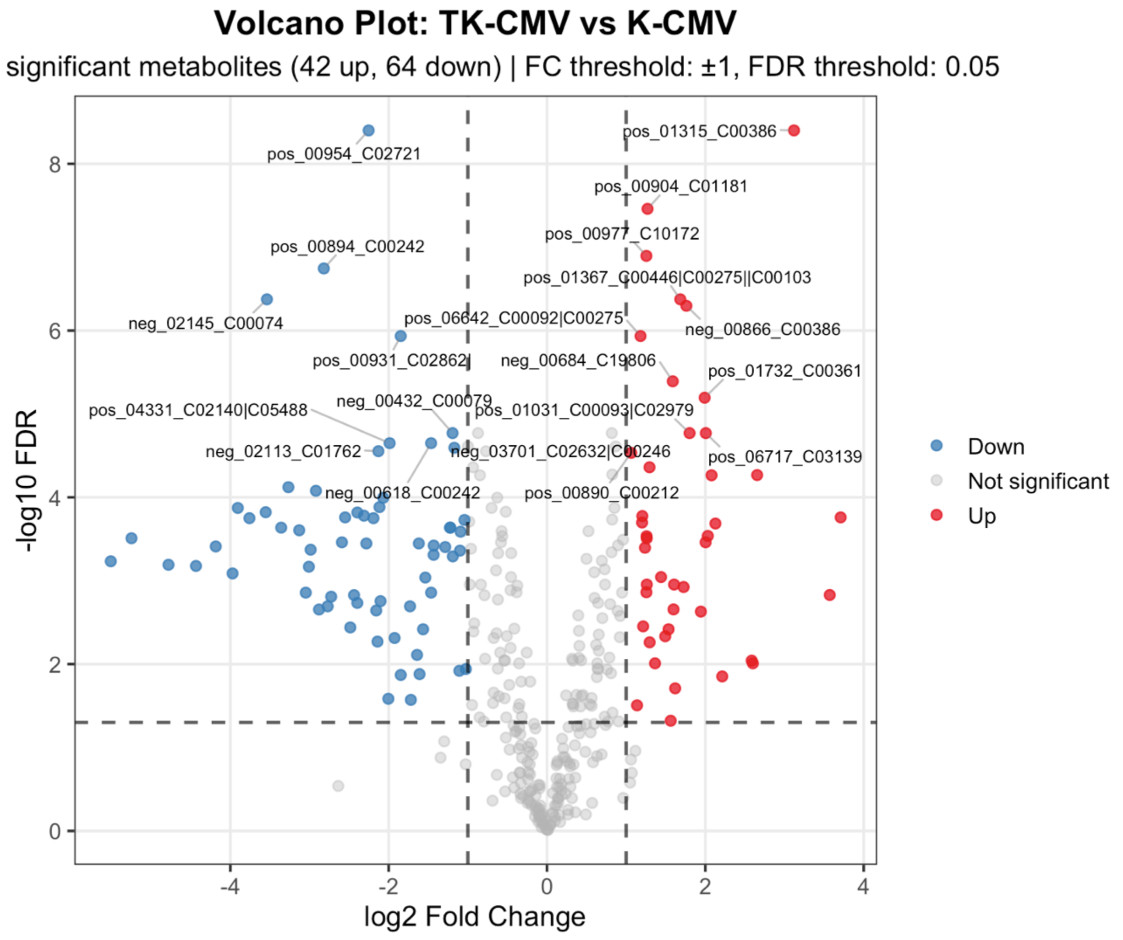


**Figure S1. Volcano plot showing differential expression between the mutant and control groups.**

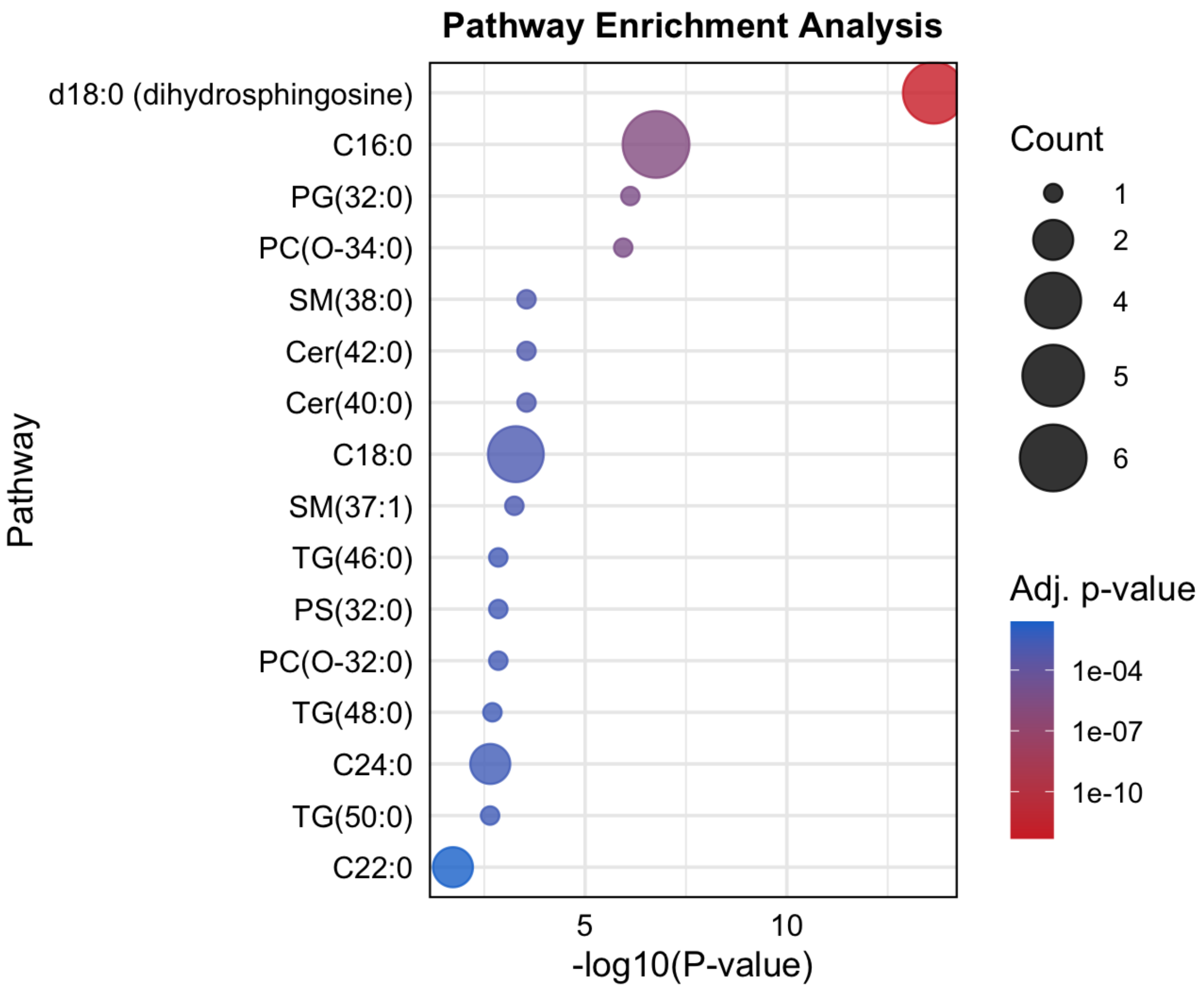


**Figure S2. Pathway enrichment analysis displaying pathways ranked by –$\log_{10}$(p-value), calculated using Fisher's exact test based on based on lipid ontology dataset derived from LION database.**
